# Prime editing is achievable in the tetraploid potato, but needs improvement

**DOI:** 10.1101/2020.06.18.159111

**Authors:** Florian Veillet, Marie-Paule Kermarrec, Laura Chauvin, Anouchka Guyon-Debast, Jean-Eric Chauvin, Jean-Luc Gallois, Fabien Nogué

## Abstract

Since its discovery and first applications for genome editing in plants, the clustered regularly interspaced short palindromic repeats (CRISPR)-Cas9 technology has revolutionized plant research and precision crop breeding. Although the classical CRISPR-Cas9 system is highly useful for the introduction of targeted small mutations for knock-out applications, this system is mostly inefficient for the introduction of precise and predictable nucleotide substitutions. Recently, the prime editing (PE) technology has been developed in human cells, allowing the introduction of all kinds of mutations, including the simultaneous generation of nucleotide transitions and transversions. Therefore, this system holds great promises for the production of gain-of-function mutants and for the improvement of precision breeding in crops. In this study, we report on the successful use of prime editing in the tetraploid and highly heterozygous potato (*Solanum tuberosum*) with the introduction of simultaneous nucleotide transitions and transversions in the *StALS1* gene.

## Introduction

Although the use of the classical CRISPR-Cas9 system brought considerable improvement for targeted mutations in plants, this system is mostly inefficient for inducing precise and predictable base modifications. If precision editing can be achieved through CRISPR-mediated gene targeting, this technology generally suffers from low efficiency due to the limited occurrence of homologous recombination in plant cells and the challenging delivery of the repair template at the cutting site (Huang and Puchta, 2019). In the last few years, several base editors have been developed, allowing cytosine and/or adenine base conversion in a generally short editing window. However, base editing suffers from some major drawbacks, such as the availability of a suitable protospacer adjacent motif (PAM) in a close vicinity of the targeted base(s), the occurrence of bystander mutations and the limited outcome diversity (Mishra et al., 2020). Recently, a new CRISPR-derived ‘search and replace’ system has been developed in human cells, named primed editing (PE), that allows the introduction of all kind of predefined mutations at a target site, including insertions, deletions and transition/transversion substitutions (Anzalone et al., 2019). Prime editors are fusion proteins made of a *Streptococcus pyogenes* Cas9 nickase (SpnCas9 H840A) and an engineered Moloney murine leukemia virus reverse transcriptase (M-MLV RT), guided to the target site by a prime editing guide RNA (pegRNA). After the ssDNA cutting about 3-bp upstream of the PAM sequence on the non-target strand, a 3’ extension of the pegRNA harboring both a primer binding site (PBS) and a reverse transcription sequence (RT sequence) allows the introduction of the desired polymorphism at the target site (Anzalone et al., 2019). As a result, this system holds great promises for precision breeding, as several traits of interest can be conferred by point mutations rather than gene loss-of-function (Henikoff and Comai, 2003). For example, point mutations in the *acetolactate synthase* (*ALS*) gene, that encodes a key enzyme in the biosynthetic pathway for branched-chain amino acids, can confer dominant resistance to ALS-inhibitors such as chlorsulfuron. Here, we report on the successful targeting of potato *StALS* genes through CRISPR-mediated plant prime editing (PPE) as a proof of concept, with the introduction of simultaneous transition and transversion mutations.

## Results and Discussion

First, we designed a pegRNA (pegRNA-StALS) with a spacer targeting *StALS1* and *StALS2* genes (Sevestre et al., 2020), and harboring a 13-bp PBS template and a 15-bp RT sequence that contains three base substitutions compared to the sequence of *StALS1* and *StALS2*, aiming at modifying both the Proline186 into a Serine and one of the three nucleotides of the PAM (synonymous mutation) to prevent nCas9 cleavage of the edited sequence (Figure 1). Because the pegRNA structure harbors an extended 3’sequence that may interfere with the Cas9 activity, we cloned the pegRNA-StALS into a modified pDeCas9 backbone (Fauser et al., 2014), resulting in the pDeCas9-StALS, for a control classical editing using *Agrobacterium*-mediated transformation with regeneration performed on kanamycin selection. Genomic DNA was extracted from regenerated potato plants and a high resolution melting (HRM) analysis was performed in order to identify mutated plants at the target locus. Eleven out of twelve regenerated transgenic plants harbored mutations at the target site (92% efficiency) (Figure 2A), which was further confirmed by Sanger sequencing (data not shown), indicating that the pegRNA structure does not impair Cas9 activity and is functional for Cas9-mediated editing.

**Figure 1:**
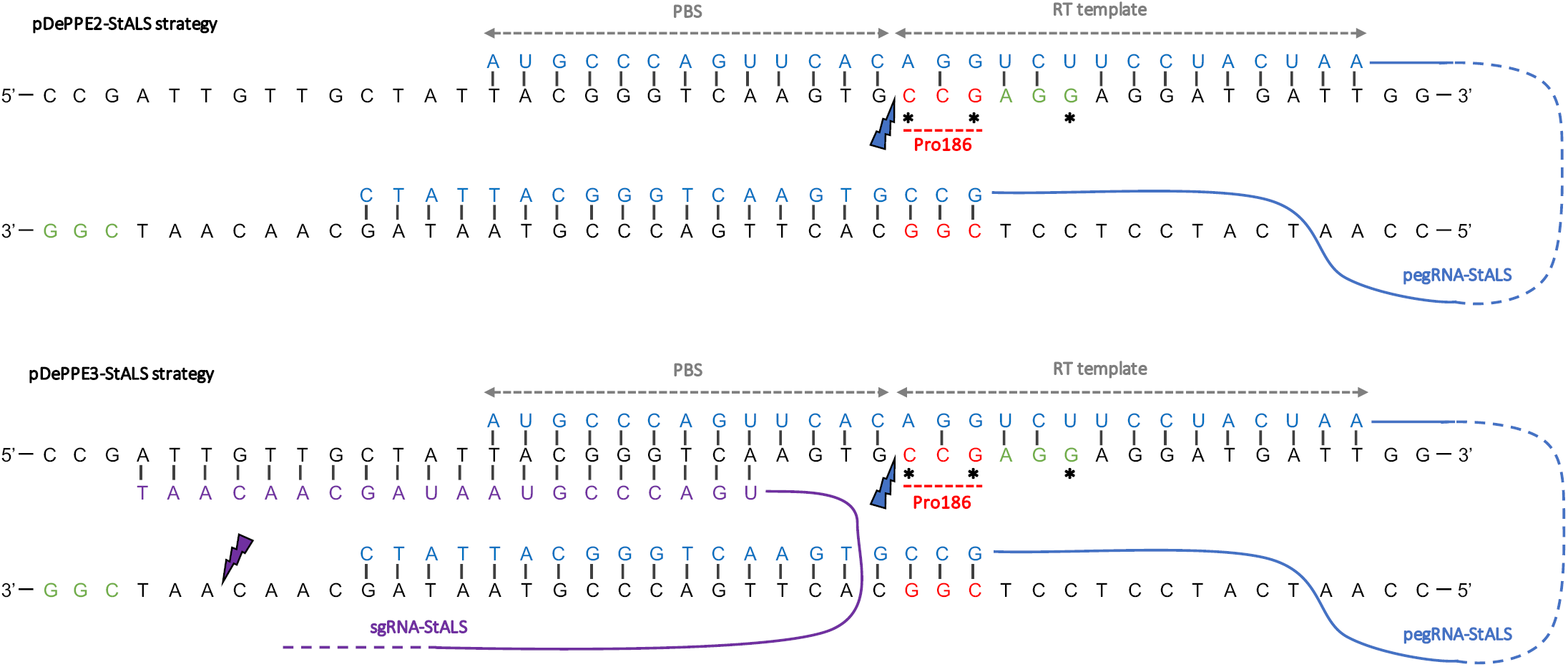
Prime editing strategy for the precise modification of the potato *ALS* genes. Schematic representation of the PPE2 and PPE3 strategies used for the modification of *StALS1* and *StALS2* genes. The PAM is represented in green, the targeted Proline in red, the targeted bases are identified with a black star and the cutting site of the pegRNA-StALS and sgRNA-StALS with a blue or purple lightning, respectively. PBS: primer binding site; RT: reverse transcriptase. The schemes are not at scale and are for illustrative purposes only.

**Figure 2:**
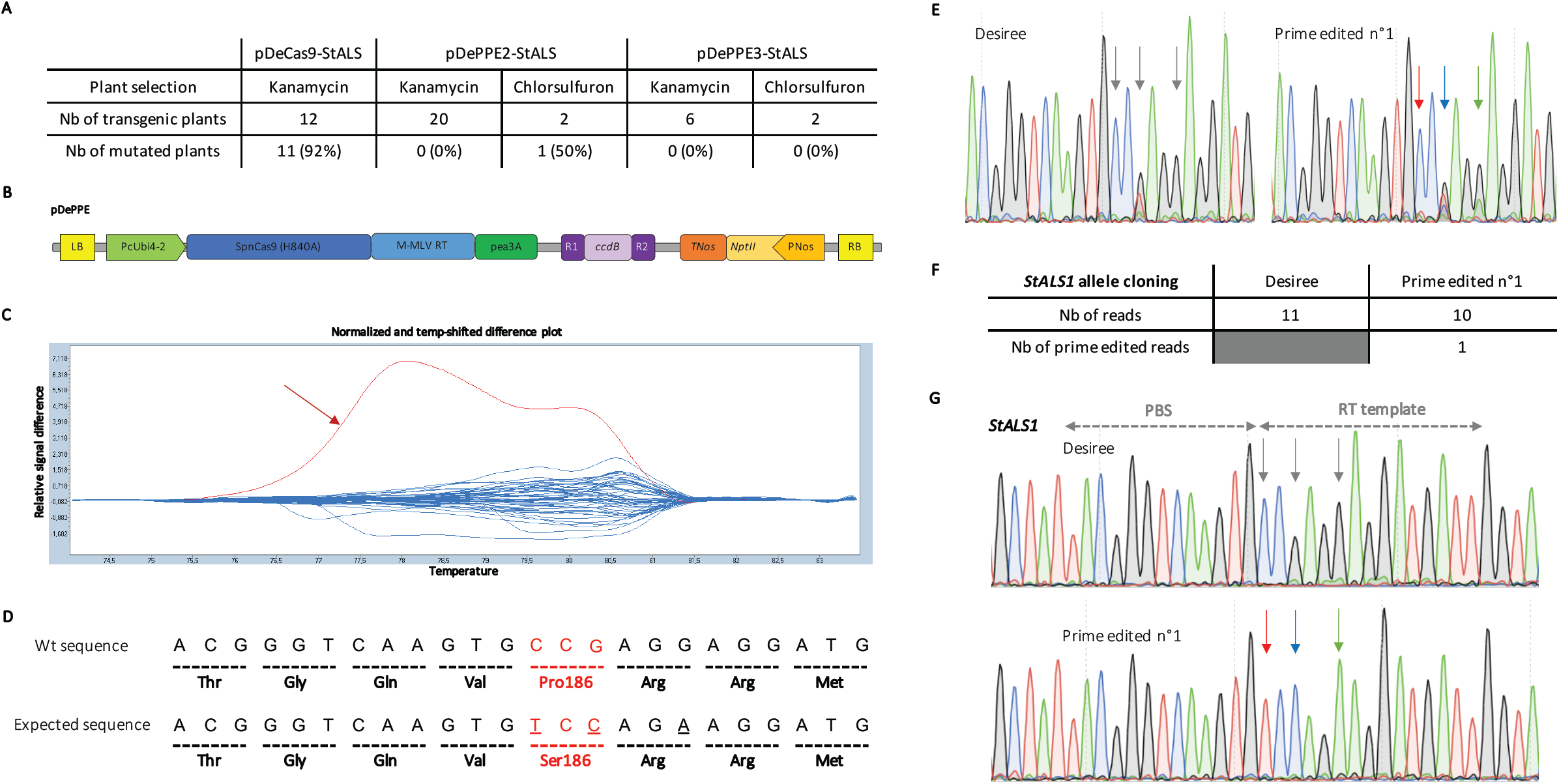
Genotyping of the prime editing targeting *StALS* loci. **A)** Summary of the molecular analysis conducted on the three conditions used in this study. **B)** Schematic representation of our prime editor for expression in dicot species through *Agrobacterium*-mediated transformation. The pDe backbone was used and the Cas9 cassette was replaced by the SpnCas9(H840A)/M-MLV RT fusion using classical cloning. The pegRNA and sgRNA were swapped with the ccdB cassette by LR gateway reaction. **C)** Screenshot of the HRM output for the pDePPE2-StALS condition is dispayed with blue curves representing wild-type profiles, while the red curve (indicated with a red arrow) represents the melting profile of a mutated plant (prime edited N°1). Both *StALS1* and *StALS2* genes were simultaneously analysed using primers matching all 8 alleles. **D)** Schematic representation of the Wt and expected sequences, with amino acids indicated under the nucleotides. The Proline 186 is in red, and targeted nucleotides are underlined. **E)** Sanger chromatograms of the targeted loci for Desiree and the prime edited plant n°1. The sequencing reaction has been performed with primers matching all the alleles of *StALS1* and *StALS2* genes. Targeted nucleotides are indicated with grey or color arrows on Desiree or mutated chromatograms, respectively. One natural SNP (G/T) is present, as evidenced by the double peak on the Desiree chromatogram. **F)** Summary of the allele cloning reaction into individual plasmids performed on Desiree and the prime edited n°1 plants after sequencing of cloned *StALS1* gene. **G)** Examples of sequencing chromatograms obtained from Desiree and prime edited n°1 plants after *StALS1* allele cloning reaction into individual plasmids. Targeted nucleotides are indicated with grey or color arrows on Wt or mutated chromatograms, respectively. A: green; T: red; C: blue; G: black.

To assess prime editing in potato, we then synthesized a dicot codon-optimized prime editor that was cloned into a modified pDe backbone, resulting into the pDePPE plasmid (Figure 2B). We then used two PPE strategies that have been demonstrated to be efficient for editing in human cells (Anzalone et al., 2019). For the PPE2 strategy, we only used the pegRNA-StALS (Figure 1). For the PPE3 strategy, we added a classical sgRNA (sgRNA-StALS) in order to cleave the non-edited strand, a method that enhanced PE efficiency in human cells (Anzalone et al., 2019) (Figure 1). These guide constructs were then cloned into the pDePPE plasmid for subsequent *Agrobacterium*-mediated transformation, resulting in the pDePPE2-StALS and pDePPE3-StALS plasmids, respectively. One week after transformation, two-thirds of the explants were transferred to a medium containing 30 ng.ml^-1^ of chlorsulfuron for the selection of edited cells, while one third of the explants were still grown on kanamycin for selection of stably transformed cells (Veillet et al., 2019b). For pDePPE2-StALS treated cells, none of the 20 kanamycin-regenerated transgenic plants were mutated at the target locus according to HRM analysis (Figure 2A). However, while one of the two plants regenerated from the chlorsulfuron-containing medium harbored a melting curve profile similar to the control, one plant displayed a mutated profile at the targeted site (Figure 2C). Sanger analyzes showed that this plant indeed harbored the 3 expected substitutions at the *StALS1* target locus (Figure 2D-E), likely on only one allele (Figure 2F-G). For the pDePPE3-StALS treated cells, none of the regenerated plants from kanamycin or chlorsulfuron-containing media was mutated according to HRM analysis (Figure 2A), suggesting that the PPE3 strategy may not substantially increase the efficiency of the system in plants.

Very recently, several reports have shown that prime editing can be applied to cereal plants (Li et al., 2020; Lin et al., 2020; Tang et al., 2020; Xu et al., 2020a; Xu et al., 2020b; Butt et al.; Hua et al.). All these studies showed that the efficiency of PPEs greatly varies between target sites and is sensitive to several parameters such as both the PBS and RT sequence. Here, we show for the first time that PPE2 strategy developed for dicot species is able to edit an endogenous gene in the cultivated tetraploid potato. Compared to our previous work on cytosine base editing in potato and *Arabidopsis* (Bastet et al., 2019; Veillet et al., 2019b; Veillet et al., 2019a; Veillet et al., 2020), the efficiency of our PPE strategy appears to be limited, as already observed for cereals. Once improved, the prime editing technology, by largely expanding the range of targeted base substitutions, could make precision breeding amenable in polyploid and vegetatively propagated crops such as potato.

## Acknowledgments

We thank Holger Puchta and his team (Botanical Institute II, Karlsruhe Institute of Technology, Karlsruhe, Germany) for providing the pDeCas9 backbone. We acknowledge the BrACySol BRC (INRAE, Ploudaniel, France) for providing the potato plants used in this study.

## References

Anzalone AV, Randolph PB, Davis JR, Sousa AA, Koblan LW, Levy JM, Chen PJ, Wilson C, Newby GA, Raguram A, et al (2019) Search-and-replace genome editing without double-strand breaks or donor DNA. Nature 576: 149–157

Bastet A, Zafirov D, Giovinazzo N, Guyon-Debast A, Nogué F, Robaglia C, Gallois J-L (2019) Mimicking natural polymorphism in *eIF4E* by CRISPR-Cas9 base editing is associated with resistance to potyviruses. Plant Biotechnol J 17: 1736–1750

Butt H, Rao GS, Sedeek K, Aman R, Kamel R, Mahfouz M Engineering herbicide resistance via prime editing in rice. Plant Biotechnology Journal. doi: 10.1111/pbi.13399

Fauser F, Schiml S, Puchta H (2014) Both CRISPR/Cas-based nucleases and nickases can be used efficiently for genome engineering in *Arabidopsis thaliana*. Plant J 79: 348–359

Henikoff S, Comai L (2003) Single-Nucleotide Mutations for Plant Functional Genomics. Annual Review of Plant Biology 54: 375–401

Hua K, Jiang Y, Tao X, Zhu J-K Precision genome engineering in rice using prime editing system. Plant Biotechnology Journal. doi: 10.1111/pbi.13395

Huang T-K, Puchta H (2019) CRISPR/Cas-mediated gene targeting in plants: finally a turn for the better for homologous recombination. Plant Cell Rep 38: 443–453

Li H, Li J, Chen J, Yan L, Xia L (2020) Precise Modifications of Both Exogenous and Endogenous Genes in Rice by Prime Editing. Molecular Plant 13: 671–674

Lin Q, Zong Y, Xue C, Wang S, Jin S, Zhu Z, Wang Y, Anzalone AV, Raguram A, Doman JL, et al (2020) Prime genome editing in rice and wheat. Nat Biotechnol 38: 582–585

Mishra R, Joshi RK, Zhao K (2020) Base editing in crops: current advances, limitations and future implications. Plant Biotechnol J 18: 20–31

Sevestre F, Facon M, Wattebled F, Szydlowski N (2020) Facilitating gene editing in potato: a Single-Nucleotide Polymorphism (SNP) map of the Solanum tuberosum L. cv. Desiree genome. Sci Rep 10: 2045

Tang X, Sretenovic S, Ren Q, Jia X, Li M, Fan T, Yin D, Xiang S, Guo Y, Liu L, et al (2020) Plant Prime Editors Enable Precise Gene Editing in Rice Cells. Molecular Plant 13: 667–670

Veillet F, Chauvin L, Kermarrec M-P, Sevestre F, Merrer M, Terret Z, Szydlowski N, Devaux P, Gallois J-L, Chauvin J-E (2019a) The Solanum tuberosum GBSSI gene: a target for assessing gene and base editing in tetraploid potato. Plant Cell Rep 38: 1065–1080

Veillet F, Perrot L, Chauvin L, Kermarrec M-P, Guyon-Debast A, Chauvin J-E, Nogué F, Mazier M (2019b) Transgene-Free Genome Editing in Tomato and Potato Plants Using Agrobacterium-Mediated Delivery of a CRISPR/Cas9 Cytidine Base Editor. IJMS 20: 402

Veillet F, Perrot L, Guyon-Debast A, Kermarrec M-P, Chauvin L, Chauvin J-E, Gallois J-L, Mazier M, Nogué F (2020) Expanding the CRISPR Toolbox in P. patens Using SpCas9-NG Variant and Application for Gene and Base Editing in Solanaceae Crops. IJMS 21: 1024

Xu R, Li J, Liu X, Shan T, Qin R, Wei P (2020a) Development of Plant Prime-Editing Systems for Precise Genome Editing. Plant Communications 1: 100043

Xu W, Zhang C, Yang Y, Zhao S, Kang G, He X, Song J, Yang J (2020b) Versatile Nucleotides Substitution in Plant Using an Improved Prime Editing System. Molecular Plant 13: 675–678

